# Adaptive Replication Fork Acceleration by CDK1-Cyclin B1 Sustains Genome Duplication despite Impaired Origin Firing

**DOI:** 10.64898/2026.05.21.726689

**Authors:** Md Shahadat Hossain, Courtney G. Sansam, Krishanu Dhar, Christopher L. Sansam

## Abstract

Although MTBP is essential for replication origin firing, we show here that strong depletion of MTBP can have minor effects on DNA replication rates. This suggests an adaptive process in the DNA replication program, so we examined mechanisms underlying this plasticity. Using an auxin-inducible degron to deplete MTBP, we found that acute suppression of MTBP blocked DNA replication, but that replication rates recovered over time. The timing of this recovery paralleled S phase expression of Cyclin B1, and inhibition of CDK1-Cyclin B1 prevented the recovery. Recovery did not involve restoration of origin firing; instead, replication recovered through accelerated fork progression. Consistent with CDK1 driving this acceleration, ATR inhibition, which activates CDK1, stimulated DNA replication in MTBP-depleted cells through CDK1-dependent increased fork progression rather than increased origin firing. Knockdown of RIF1, a known CDK1 target, phenocopied this effect. Although RIF1 is best known for opposing DDK-dependent MCM phosphorylation at origins, we find that RIF1 knockdown stimulates replication even when DDK is inhibited. Furthermore, RIF1 loss increased replication by accelerating fork progression rather than increasing origin firing. Together, these findings reveal a CDK1-RIF1-dependent mechanism that promotes fork speed during S phase and defines a form of replication plasticity in which fork rate compensates for reduced origin firing.

**SIGNIFICANCE STATEMENT:** Accurate genome duplication requires thousands of replication origins to fire and replication forks to complete DNA synthesis on schedule. When origin firing is compromised, it is unclear how cells avoid replication failure. We show that cells adapt to persistent loss of the origin-firing factor MTBP by accelerating replication fork progression through a CDK1–RIF1-dependent mechanism, partially compensating for reduced initiation. This adaptive response defines a form of replication plasticity in which cells rebalance origin usage and fork speed to sustain DNA synthesis. This mechanism may be especially relevant in cancer cells or other contexts where replication initiation is chronically stressed.

## INTRODUCTION

Faithful genome duplication requires coordinated activation of thousands of replication origins and orderly progression of replication forks to complete S phase on schedule (1, 2). In eukaryotic cells, replication initiation begins when ORC, CDC6, and CDT1 load MCM2-7 helicase complexes at origins as inactive double hexamer (3-6). DDK then phosphorylates loaded MCM complexes to promote recruitment of firing factors and prime origins for activation (7-12). In vertebrates, S-phase CDK promotes CMG (CDC45 - MCM2-7 - GINS) assembly through firing factors MTBP, TRESLIN/TICRR, and DONSON (13-23). Once assembled, CMG helicases unwind DNA and establish bidirectional replication forks that support genome duplication (11, 24-26).

Cyclin-dependent kinases are central regulators of this process, as they coordinate replication initiation with cell-cycle progression. During an unperturbed cell cycle, CDK2, in complex with Cyclin E or Cyclin A, is widely regarded as a major driver of S-phase entry and replication origin firing (27, 28). In contrast, CDK1-Cyclin B activity is generally restrained during S phase, increasing as mitotic-entry network activity rises around the S/G2-to-mitosis transition in order to preserve the temporal separation of DNA replication and mitosis (29-33). This canonical division of function places CDK2 at the center of replication control, largely excluding CDK1 from S-phase regulation.

Intriguingly, accumulating evidence indicates that CDK1 can promote DNA replication under specific conditions (34-37). CDK1 can compensate for CDK2 loss and drive the G1-to-S phase transition in CDK2-deficient cells, demonstrating functional redundancy within the CDK network (36-39). CDK1 has also been shown to substitute for CDC7 when CDC7 is inhibited during the G1/S transition (34). During an unperturbed S phase, CDK1 appears to play only a minor role in origin firing. However, CDK1 can be activated to stimulate replication when the WEE1 or ATR checkpoint kinases that serve to restrain initiation, are inhibited (40-45). Despite these observations, the contribution of CDK1 to DNA synthesis, particularly when canonical initiation pathways are compromised, remains incompletely understood.

The overall rate of DNA synthesis reflects the combined contributions of replication origin usage and replication fork progression (46, 47). In multiple systems, including mammalian cells and yeast, high origin density is often associated with slower fork movement, whereas reduced origin firing can permit faster fork progression (48-50). This inverse relationship is thought to arise from competition for limiting replication factors and nucleotides, as well as from checkpoint signaling elicited by increased numbers of active forks (48, 50, 51). For example, embryonic stem cells maintain high origin density and relatively slow but stable fork progression, whereas differentiated cells exhibit early S-phase fork slowing followed by ATR-dependent fork acceleration later in S phase (52). How checkpoint pathways dynamically balance origin usage and fork speed to sustain DNA synthesis when initiation is impaired remains an open question.

Here, we examine how cells adapt to compromised MTBP-dependent origin firing. We find that MTBP depletion initially suppresses DNA synthesis, but the overall synthesis rates subsequently recover toward normal, despite persistently low levels of origin firing and reduced CDC45 loading. This recovery is driven by CDK1-Cyclin B-dependent acceleration of replication fork progression rather than by restoration of initiation. We further show that this adaptive increase in fork speed is restrained by ATR signaling, and we identify RIF1 as a downstream effector of CDK1 in this response. Our data support a model in which CDK1 promotes replication fork acceleration, at least in part by phosphorylating RIF1; this acceleration partially compensates for impaired origin firing. Together, these findings define a CDK-centered regulatory switch that enables cells to rebalance origin usage and fork progression to sustain genome duplication when initiation is compromised. This mechanism of replication plasticity may be especially relevant in stressed or highly proliferative cells.

## RESULTS

### DNA synthesis recovers over time during sustained MTBP depletion

MTBP is a canonical origin-firing factor and is considered rate-limiting for replication initiation. Our findings, however, indicate that the consequences of its knockdown vary across cell lines. In particular, we have found that HCT116 cells replicate DNA at relatively normal rates, despite MTBP knockdown (41). We wondered whether this might result from adaptation to MTBP loss over time. To determine how DNA synthesis responds to sustained MTBP depletion, we used an HCT116 line in which endogenous MTBP is fused to mClover and a miniAID degron (hereafter MTBP-AID)(41, 53, 54). To compare the effects of immediate (2 hours) and prolonged (20 hours) MTBP depletion, we treated MTBP-AID cells with auxin (IAA) and pulse-labeled with EdU for 30 minutes; DNA synthesis (EdU incorporation) and DNA content were quantified by flow cytometry (Fig. 1B). Two hours of auxin treatment reduced EdU incorporation throughout S phase relative to untreated controls, consistent with the established requirement for MTBP in replication origin firing (Fig. 1A,C). In contrast, with prolonged auxin treatment, EdU incorporation in early and mid S phase cells progressively recovered, approaching near-control levels by 16-20 hours despite continued MTBP depletion (Fig. 1A,C). Replication in late S phase cells, however, remained impaired (Fig. 1A,C,D). In parallel, a population of late S-phase cells with low EdU incorporation emerged and gradually accumulated (Fig. 1D), consistent with incomplete replication of late-replicating regions. The G1 fraction also transiently increased approximately 1.5-fold between 2 and 8 hours of auxin treatment before returning toward baseline by 16 hours (Fig. 1D). To confirm that these effects reflected sustained MTBP loss, we measured chromatin-bound and total MTBP by anti-GFP staining of the mClover tag. MTBP dropped to near-background and remained depleted throughout the 20-hour time course (Fig. 1E-H). Together, these findings indicate that HCT116 cells can adapt to sustained impairment of origin firing through MTBP-independent compensatory mechanisms.

**Figure 1.**
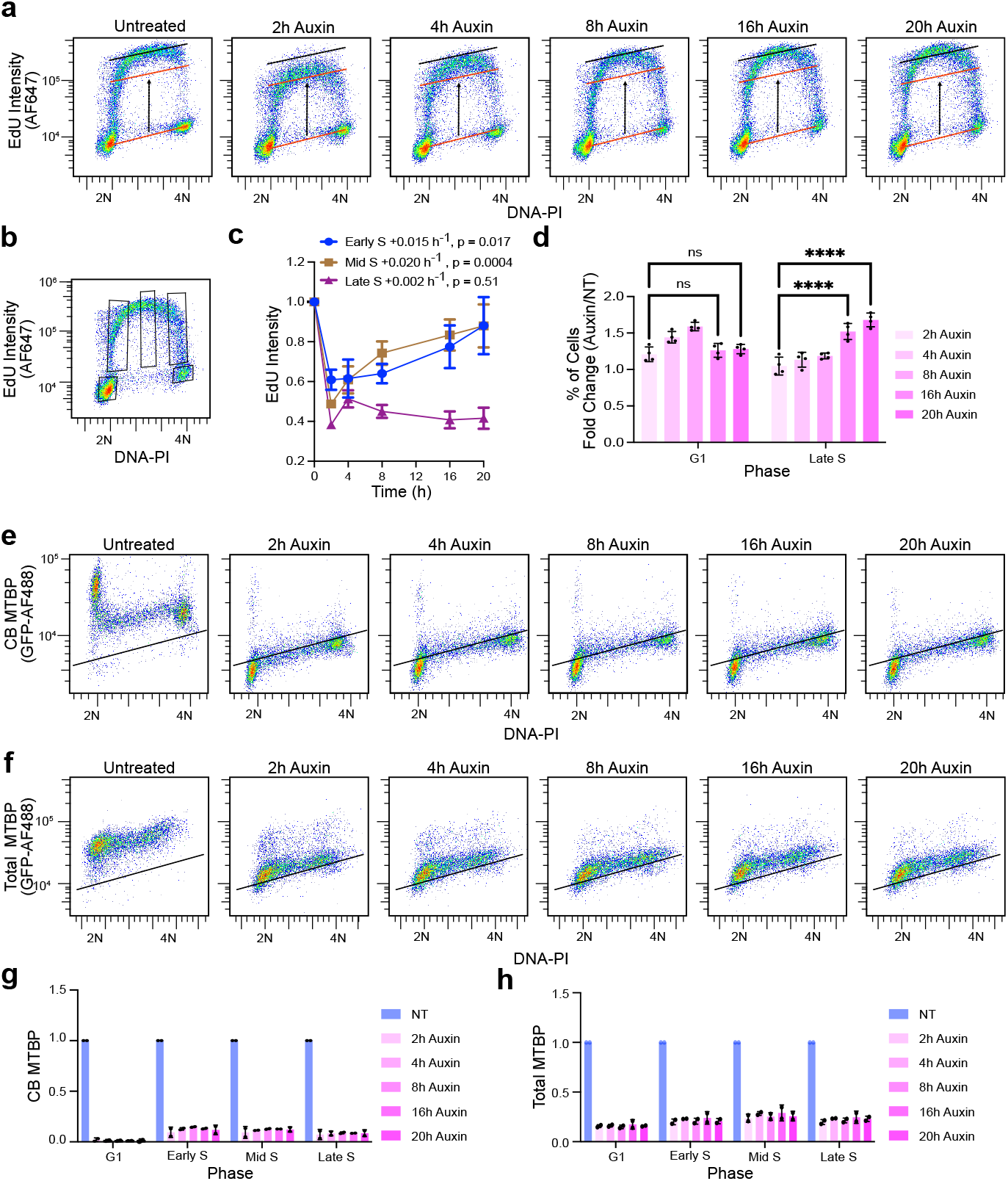
DNA synthesis adapts to sustained loss of MTBP. (a) Recovery of DNA synthesis during auxin-induced MTBP depletion, measured by flow cytometry. Pseudocolor plots show EdU incorporation (y-axis, log scale) versus DNA content (propidium iodide; x-axis, linear scale). Lines and arrows indicating EdU incorporation levels were drawn on the untreated plot and overlaid on all other plots to aid visual comparison. (b) Gating strategy used to define early, mid, and late S phase. Gates were defined on the untreated sample and applied uniformly across all time points. (c) Quantification of (a). Median EdU signal was normalized to G1 within each sample and then to the untreated condition within each subphase of S. Line plots represent mean ± SEM from four independent experiments. Statistical comparisons by linear regression. (d) G1 and late S phase percentages following MTBP depletion. Values represent the ratio of auxin-treated to untreated samples at each time point. Bars represent mean ± SD from four independent experiments. Statistical comparisons by two-way ANOVA with Tukey’s multiple comparisons test. (e) Chromatin-bound MTBP (CB-MTBP; anti-GFP) levels during auxin-induced depletion, measured by flow cytometry. Pseudocolor plots as in (a). (f) Total MTBP levels during auxin-induced depletion, measured by flow cytometry. Pseudocolor plots as in (a). (g) Quantification of (e), normalized to the untreated condition within each cell-cycle gate. Bars represent mean ± SD from two independent experiments. (h) Quantification of (f), normalized to the untreated condition within each cell-cycle gate. Bars represent mean ± SD from two independent experiments.

### CDK1-Cyclin B activity contributes to recovery of DNA synthesis after MTBP depletion

Previous studies showed that MTBP depletion increases Cyclin B1 (CCNB1) expression during S phase (55), suggesting that elevated CDK1-CCNB1 activity may promote replication in MTBP-deficient cells. To test this hypothesis, we measured nuclear CCNB1 levels as a function of DNA content during an auxin treatment time course in MTBP-AID HCT116 cells. Flow cytometric analysis revealed a progressive increase in nuclear CCNB1 across all cell-cycle stages, including early, mid, and late S phase, following auxin addition (Fig. 2A, B). We next assessed total CCNB1 levels after 20 hours of auxin treatment and found that they also increased, consistent with the rise in nuclear CCNB1 (Fig. 2C, D). This accumulation paralleled the gradual recovery of DNA synthesis observed in Figure 1, suggesting that elevated CCNB1 levels may contribute to the restoration of replication during prolonged MTBP depletion.

**Figure 2.**
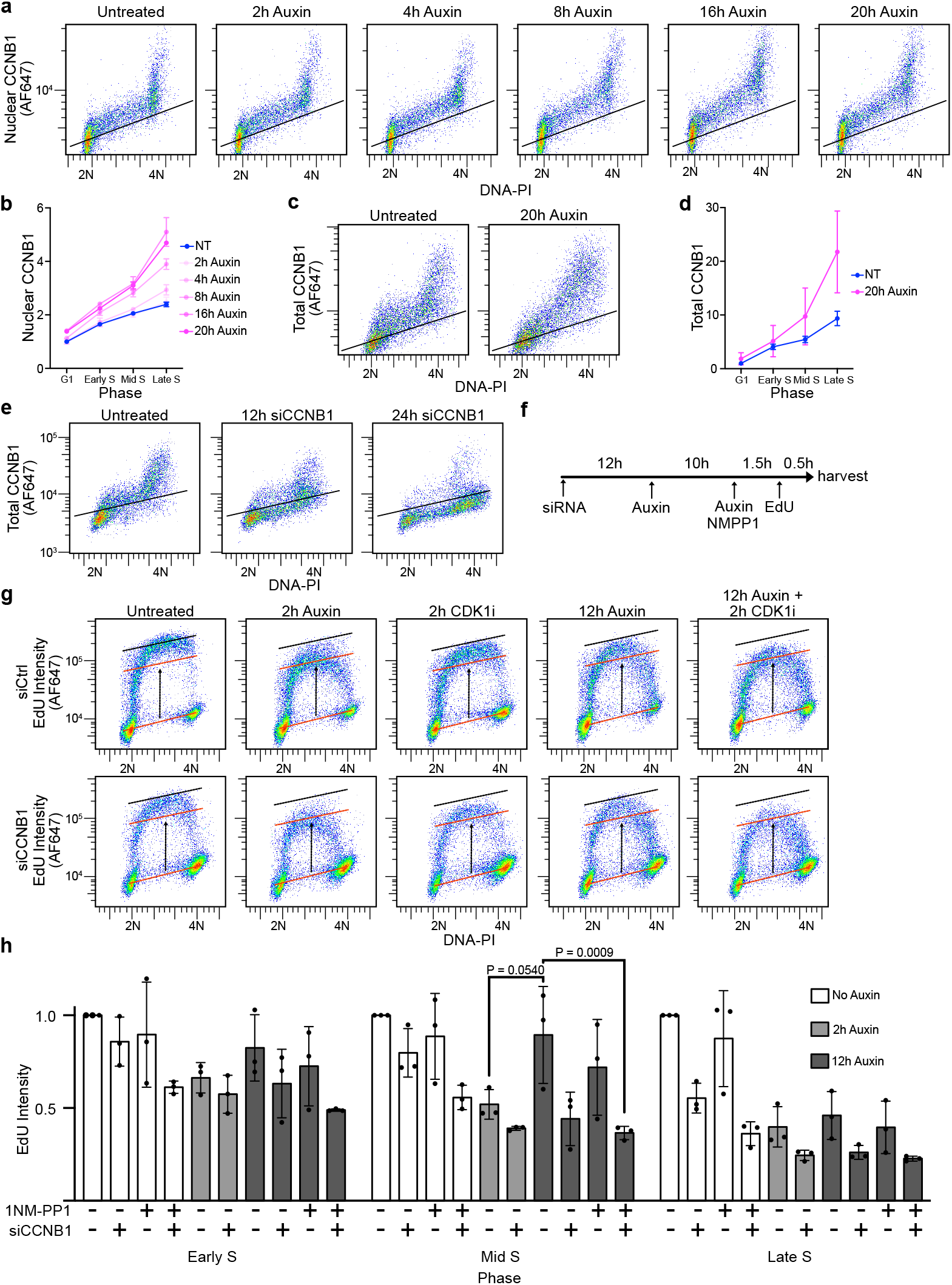
Nuclear CCNB1-CDK1 activity contributes to recovery of DNA synthesis following MTBP depletion. (a) Nuclear CCNB1 levels across the cell cycle following auxin-induced MTBP depletion, measured by flow cytometry. Pseudocolor plots show nuclear CCNB1 intensity (y-axis, log scale) versus DNA content (propidium iodide; x-axis, linear scale). Black line marks background (no primary antibody) signal. (b) Quantification of (a). Median CCNB1 signal was normalized to the untreated condition within each cell-cycle phase. Line plots represent mean ± SEM from two independent experiments. (c) Effect of prolonged MTBP depletion on total CCNB1 levels, measured by flow cytometry. Pseudocolor plots as in (a). (d) Quantification of (c), normalized to the untreated condition. Line plots represent mean ± SEM from three independent experiments. (e) Validation of siCCNB1 knockdown efficiency, measured by flow cytometry. Pseudocolor plots as in (a). (f) Experimental timeline for siRNA transfection and drug treatments in (g) and (h). (g) Effect of CCNB1 siRNA knockdown and CDK1 inhibition on recovery of DNA synthesis following MTBP depletion, measured by flow cytometry. Pseudocolor plots as in (a). Lines and arrows indicating EdU incorporation levels were drawn on the siControl untreated plot and overlaid on all other plots to aid visual comparison. (h) Quantification of (g), normalized to the siControl untreated condition within each subphase of S. Bars represent mean ± SD.

To determine whether CDK1-CCNB1 activity contributes causally to this recovery, we generated an analog-sensitive CDK1 cell line in which endogenous CDK1 was replaced with a *Xenopus* CDK1^AS^ transgene, allowing for acute inhibition with the ATP analog 1NM-PP1 (hereafter MTBP-AID-CDK1^AS^)(33, 41). In parallel, *CCNB1* was depleted by siRNA and knockdown was confirmed by flow cytometry (Fig. 2E). We then measured DNA synthesis in MTBP-AID-CDK1^AS^ cells treated with auxin and/or 1NM-PP1 for 2 or 12 hours, with or without CCNB1 knockdown (Fig. 2F). As in Figure 1, 2 hours of auxin inhibited EdU incorporation, whereas 12 hours partially restored synthesis; this recovery was blocked by combined CDK1 inhibition and CCNB1 depletion (Fig. 2G, H). Together, these results indicate that the rise in CDK1-CCNB1 activity following MTBP loss promotes the recovery of DNA synthesis.

### ATR limits DNA synthesis during MTBP depletion in a CDK1-dependent manner

The ATR-CHK1-CDC25 pathway restrains DNA synthesis in unperturbed cells, primarily by limiting origin firing through inhibition of CDK1 and CDK2 (44, 45, 56, 57). Our finding that DNA synthesis recovers during MTBP depletion, however, suggested that CDK1 may promote replication through an additional mechanism not explained by increased origin firing. We therefore asked whether ATR signaling also restrains this alternative CDK1-dependent replication mechanism. To test this, we treated MTBP-AID-CDK1^AS^ cells with an ATR inhibitor, with or without MTBP knockdown and/or CDK1 inhibitor, and measured DNA synthesis by flow-cytometric analysis of EdU incorporation. If ATR limits this alternative CDK1-dependent mechanism, then ATR inhibition should stimulate DNA synthesis in MTBP-depleted cells, and that effect should be blocked by CDK1 inhibition.

Consistent with the established role of ATR in suppressing DNA replication during S phase, ATR inhibitor treatment for 4 hours increased EdU incorporation in control cells (Fig. 3A-D). ATR inhibition also stimulated EdU incorporation in cells where DNA synthesis had been reduced by MTBP depletion, whether by 8 hours of auxin treatment or 48 hours of siMTBP (Fig. 3A-D). These results show that ATR continues to restrain DNA synthesis even when MTBP-dependent initiation is compromised.

**Figure 3.**
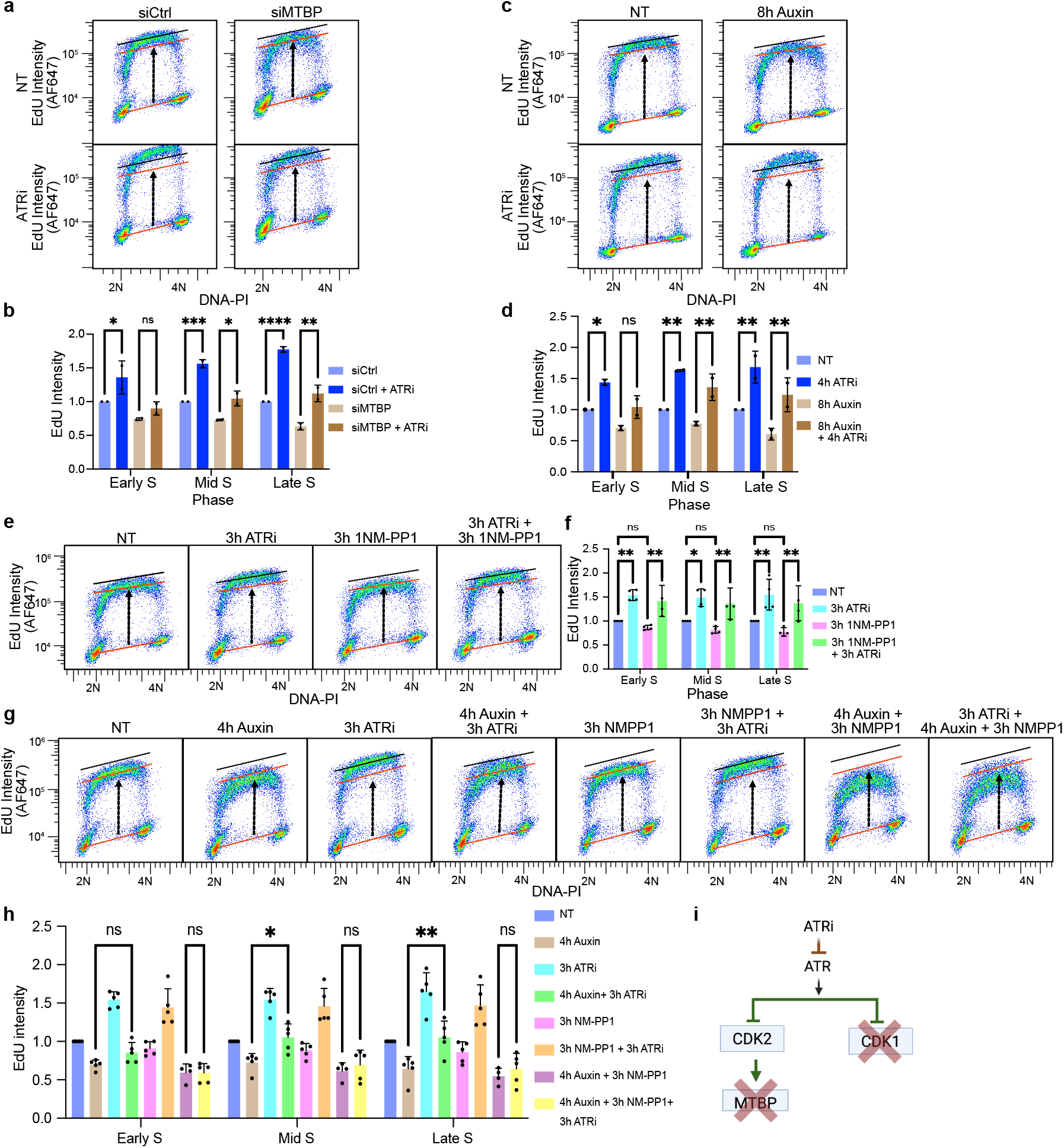
ATR limits DNA synthesis during MTBP depletion in a CDK1-dependent manner. (a) Effect of ATR inhibition on DNA synthesis in siRNA-transfected cells, measured by flow cytometry. Pseudocolor plots show EdU incorporation (y-axis, log scale) versus DNA content (propidium iodide; x-axis, linear scale). Lines and arrows indicating EdU incorporation levels were drawn on the siControl NT plot and overlaid on all other plots to aid visual comparison. NT, untreated. (b) Quantification of (a). Median EdU signal was normalized to the siControl NT condition within each subphase of S. Bars represent mean ± SD. Statistical comparisons by two-way ANOVA with Tukey’s multiple comparisons test. (c) Effect of ATR inhibition on DNA synthesis in MTBP-AID cells after auxin-induced MTBP depletion, measured by flow cytometry. Pseudocolor plots as in (a). (d) Quantification of (c), normalized to the NT condition within each subphase of S. Bars represent mean ± SD. Statistical comparisons as in (b). (e) Effect of CDK1 inhibition, alone or in combination with ATR inhibition, on DNA synthesis without MTBP depletion, measured by flow cytometry. Pseudocolor plots as in (a). (f) Quantification of (e), normalized to the NT condition within each subphase of S. Bars represent mean ± SD from four independent experiments. Statistical comparisons as in (b). (g) Effect of CDK1 inhibition on ATR inhibitor-stimulated DNA synthesis in MTBP-depleted cells, measured by flow cytometry. Pseudocolor plots as in (a). (h) Quantification of (g), normalized to the NT condition within each subphase of S. Bars represent mean ± SD. Statistical comparisons as in (b). (i) Model: ATR suppresses both MTBP-dependent and CDK1-dependent pathways of DNA replication stimulation.

We next tested whether ATR-regulated DNA synthesis depends on CDK1. In control cells, ATR inhibitor treatment for 3 hours increased DNA synthesis, consistent with relief of ATR-CHK1-mediated inhibition of CDK activity (Fig. 3E, F). 1NM-PP1 treatment in the MTBP-AID-CDK1^AS^ cells alone caused a slight decrease in EdU incorporation. Importantly, ATR inhibition increased DNA synthesis to a similar extent with or without 1NM-PP1, indicating that ATR-stimulated replication does not require CDK1 activity in the absence of MTBP depletion (Fig. 3E, F).

To determine whether this CDK1 independence holds in MTBP-depleted cells, we tested the effect of CDK1 inhibition, MTBP depletion, or their combination on ATRi-stimulated DNA synthesis (Fig. 3G, H). MTBP-AID-CDK1^AS^ cells were treated with auxin for 4 hours to deplete MTBP and then exposed to 1NM-PP1 and/or ATR inhibitor. Auxin treatment alone reduced DNA synthesis; ATR inhibition increased DNA synthesis; 1NM-PP1 alone had minimal effect. However, combined CDK1 inhibition and MTBP depletion caused a pronounced reduction in EdU incorporation, and under these conditions ATR inhibition no longer stimulated replication. These results demonstrate that CDK1 activity becomes essential for ATRi-induced DNA synthesis when MTBP is depleted. Together, these data indicate that MTBP loss rewires replication control such that ATR-mediated stimulation of DNA synthesis becomes CDK1-dependent, likely reflecting an adaptive shift toward CDK1-CCNB1-driven replication (Fig. 3I).

### Replication persists despite reduced CDC45 loading after MTBP depletion

To investigate how ATR and CDK1 regulate DNA synthesis when MTBP depletion inhibits origin firing, we asked whether these conditions alter CDC45 loading onto chromatin. Because CDC45 is an essential component of the active replicative helicase, its abundance on chromatin provides a readout of replication fork activation. To measure CDC45 directly, we generated an HCT116 cell line in which endogenous CDC45 was tagged with mClover, allowing us to quantify CB-CDC45 across the cell cycle by anti-GFP flow cytometry. In parallel, because our earlier results implicated CDK1 in sustaining DNA synthesis after MTBP depletion, we used an analog-sensitive CDK1 (CDK1^AS^) background so that 1NM-PP1 could acutely inhibit CDK1 activity (hereafter CDC45-mClover-CDK1^AS^)(Fig. 4C, D). These tools allowed us to test whether ATR inhibition stimulates CDC45 loading, and whether any such effect requires MTBP and/or CDK1 activity.

**Figure 4.**
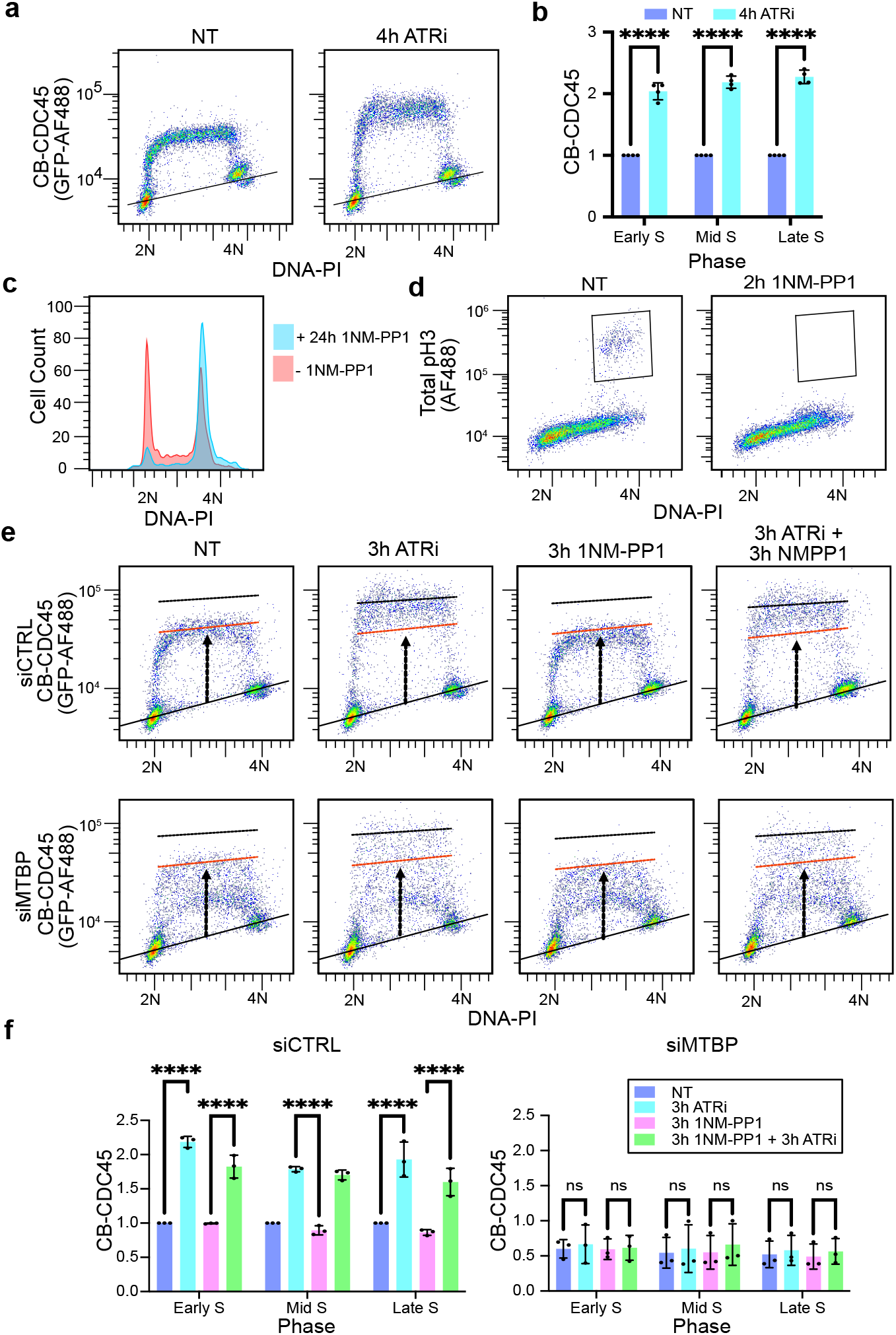
ATR restricts CDC45 loading through an MTBP-dependent, CDK1-independent mechanism. (a) Effect of ATR inhibition on chromatin-bound CDC45 (CB-CDC45) levels, measured by flow cytometry. Pseudocolor plots show CB-CDC45 signal (y-axis, log scale) versus DNA content (propidium iodide; x-axis, linear scale). Lines and arrows indicating CB-CDC45 levels were drawn on the NT plot and overlaid on all other plots to aid visual comparison. NT, untreated. (b) Quantification of (a), normalized to the untreated condition within each cell-cycle gate. Bars represent mean ± SD from four independent experiments. Statistical comparisons by two-way ANOVA with Tukey’s multiple comparisons test. (c) Cell-cycle distribution following CDK1 inhibition, measured by flow cytometry. Histograms show cell count versus DNA content. (d) Confirmation of mitotic entry inhibition by CDK1 inhibitor, assessed by phospho-histone H3 staining versus DNA content, measured by flow cytometry. (e) Effect of ATR inhibition and CDK1 inhibition on CB-CDC45 levels in MTBP-knockdown cells, measured by flow cytometry. Pseudocolor plots as in (a). (f) Quantification of (e), normalized to the siControl untreated condition within each cell-cycle gate. Bars represent mean ± SD from three independent experiments. Statistical comparisons as in (b).

In untreated CDC45-mClover-CDK1^AS^ cells, relative to G1 and G2/M, CB-CDC45 increased across S phase as expected, and ATR inhibitor treatment for 4 hours further elevated CB-CDC45 throughout S phase (Fig. 4A, B). CB-CDC45 was strongly reduced 48 hours after siMTBP transfection; ATR inhibition increased CB-CDC45 in siControl cells but failed to do so in siMTBP-treated cells (Fig. 4E, F). CDK1 inhibition had little effect on CB-CDC45 with or without ATR inhibition. Thus, ATR inhibition promotes CDC45 loading in an MTBP-dependent, CDK1-independent manner, indicating that ATRi-stimulated DNA synthesis in MTBP-depleted cells is unlikely to reflect increased origin firing.

### Replication fork progression accelerates when replication initiation is suppressed

Our finding that CDC45 loading depends on MTBP suggests that CDK1-dependent DNA synthesis, whether adaptive or ATR inhibitor-stimulated, is unlikely to reflect increased origin firing. A known inverse relationship between fork number and fork speed predicts an alternative mechanism: when fewer origins fire, existing forks elongate faster. Consistent with this, published studies have shown that siRNA-mediated MTBP depletion reduces fork number while increasing fork speed in U2OS cells (58, 59). We therefore asked whether fork acceleration also drives the recovery of DNA synthesis following prolonged MTBP depletion in our HCT-116 auxin-degron system. To address this, we used nanopore sequencing to measure EdU and BrdU incorporation following 20 hours of auxin-induced MTBP depletion (Fig. 5A). MTBP-AID cells were pulse-labeled sequentially with EdU (10 min) and BrdU (10 min), followed by a 40-minute thymidine chase. DNAscentV2 was then applied to each nanopore sequencing read to classify replication events as initiations, terminations, or elongating forks based on the spatial pattern of EdU and BrdU incorporation along individual DNA molecules (60, 61). Both initiation and termination events were markedly reduced after 20 hours of auxin treatment (Fig. 5B). In contrast, replication track length increased, indicating that recovery of DNA synthesis under these conditions is driven by increased fork elongation rather than initiation (Fig. 5C).

**Figure 5.**
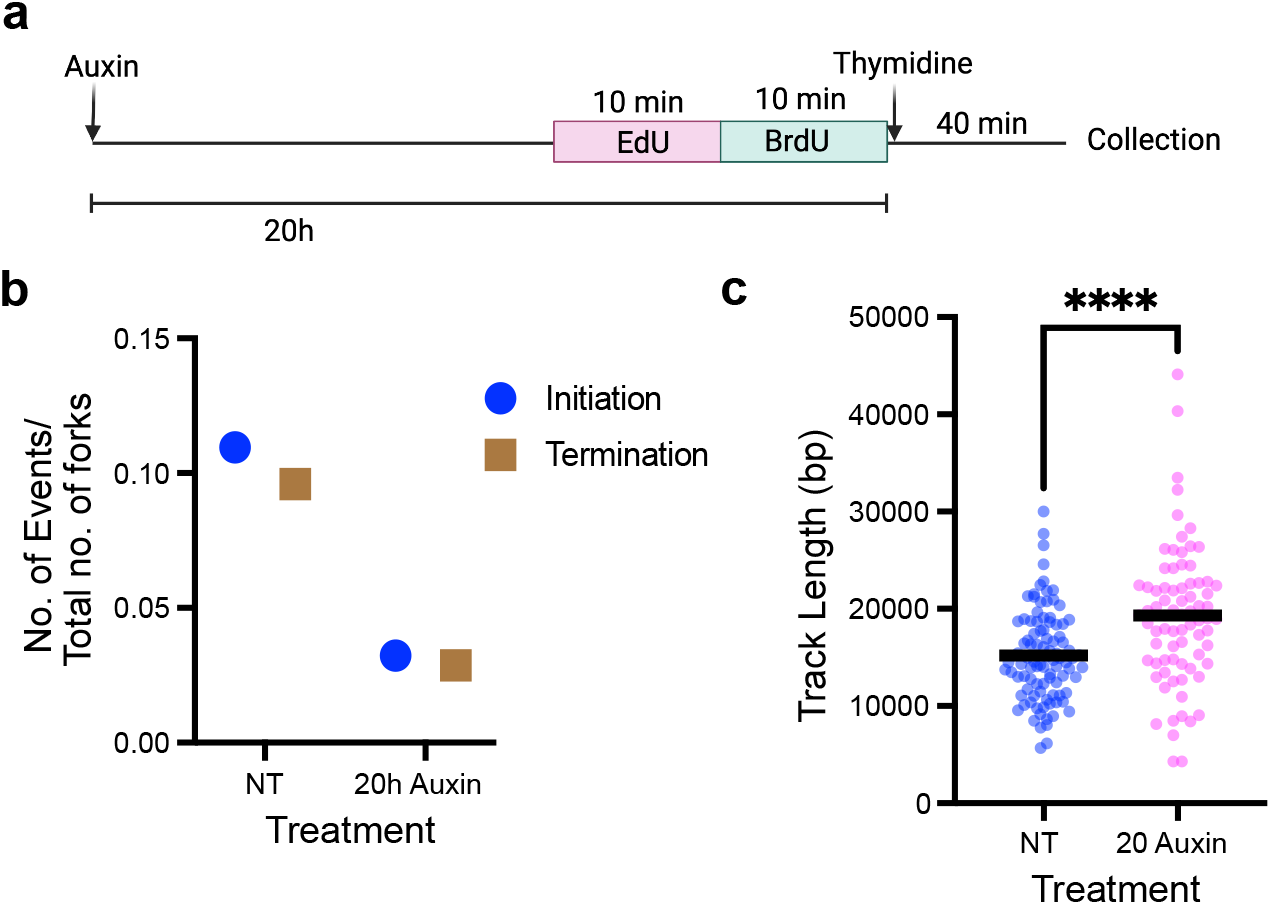
Nanopore analysis confirms reduced initiation but increased fork elongation following MTBP depletion. (a) Schematic of the labeling and treatment scheme. Cells were treated with auxin for 20 hours. During the final 20 minutes, cells were sequentially labeled with EdU (10 min) and BrdU (10 min), followed by a thymidine chase (40 min) before collection. (b) Quantification of initiation and termination events, shown as number of events normalized to total forks. Y axis: events per total forks; X axis: treatments (NT, 20 h auxin). (c) Replication track length under indicated conditions. Y axis: track length; X axis: treatments (NT, 20 h auxin).

### RIF1 acts downstream of CDK1 to restrain replication fork progression

Because CDK1–Cyclin B1 activity is required for recovery of DNA synthesis in MTBP-depleted cells (Fig. 2G, H), and because CDK1 has been shown to directly phosphorylate and inhibit RIF1, we tested whether RIF1 acts downstream of CDK1 to promote replication (44, 45). If RIF1 were the relevant downstream target, RIF1 depletion should phenocopy MTBP loss by increasing DNA synthesis. To test this prediction, we used HCT116 cells in which endogenous RIF1 is fused to mClover and an auxin-inducible degron and CDK1 was replaced with the analog-sensitive mutant (Fig. 6A-E; hereafter RIF1-AID-CDK1^AS^)(62). RIF1 depletion in RIF1-AID-CDK1^AS^ cells increased EdU incorporation in mid and late S phase, confirming the function of RIF1 as a DNA replication inhibitor (63, 64). By contrast, CDK1 inhibition with 1NM-PP1 decreased EdU incorporation, suggesting that low-level CDK1 activity modestly promotes replication in unperturbed cells. RIF1 depletion stimulated replication even in 1NM-PP1-treated cells (Fig. 6F, G). Thus, RIF1 depletion bypasses the reduction in DNA synthesis caused by CDK1 inhibition, supporting the model that RIF1 acts downstream of CDK1 in regulating replication.

**Figure 6.**
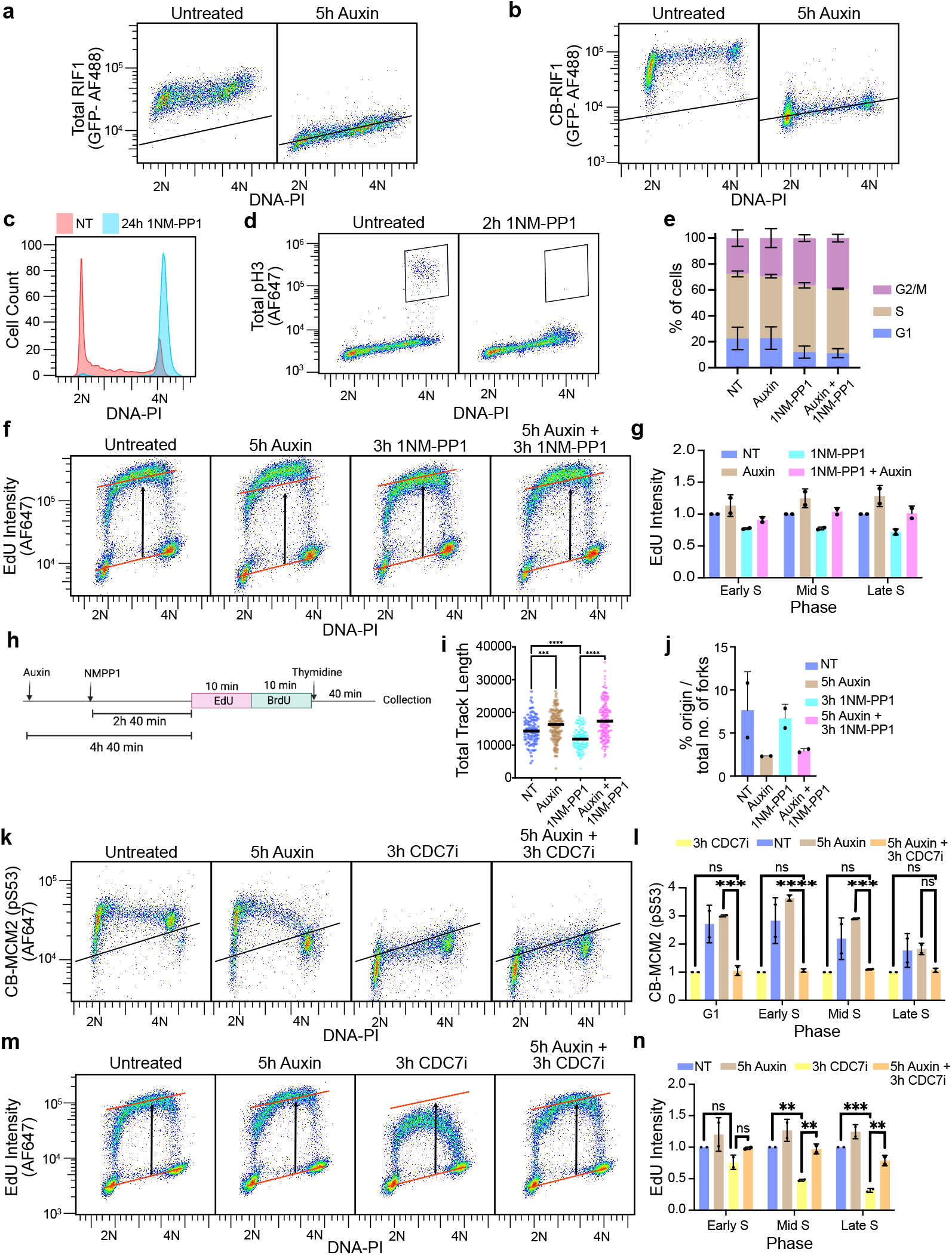
CDK1-dependent recovery of DNA synthesis occurs through inhibition of RIF1 and enhanced fork progression rather than increased origin firing. (a) Flow cytometry analysis of total RIF1 levels in untreated cells and cells treated with auxin for 5 hours. RIF1 was detected using anti-GFP antibody recognizing the mClover tag. (b) Chromatin-bound RIF1 (CB-RIF1) under the same conditions as in (a); cells were extracted to isolate insoluble nuclear fractions prior to anti-GFP staining. Pseudocolor plots in (a) and (b) show the indicated signal (y-axis, log scale) versus DNA content (propidium iodide, x-axis, linear scale). (c) DNA content distribution in untreated cells (NT; red) or cells treated with CDK1 inhibitor (1NM-PP1; blue) for 24 hours. (d) Phospho-histone H3 (pH3) versus DNA content in untreated or 1NM-PP1-treated cells, confirming inhibition of mitotic entry. (e) Cell-cycle distribution under the indicated conditions. Bars represent the percentage of cells in G1, S, and G2/M. (f) EdU incorporation versus DNA content in cells treated with auxin, 1NM-PP1, or both. Pseudocolor plots as in (a). (g) Quantification of (f). Median EdU signal was normalized to the untreated condition within each S-phase fraction. Bars represent mean ± SD from two independent experiments. (h) Schematic of the nanopore sequencing experimental design. Cells were treated with auxin for 4 h 40 min and 1NM-PP1 for 2 h 40 min, then sequentially pulse-labeled with EdU (10 min) and BrdU (10 min) in the continued presence of drugs, followed by a 40-min thymidine chase before collection. (i) Total replication track length under the indicated conditions, measured by nanopore sequencing. (j) Origin firing, expressed as the percentage of initiation events relative to total forks, under untreated, 5 h auxin, 3 h 1NM-PP1, or 5 h auxin plus 3 h 1NM-PP1 conditions. (k) Chromatin-bound MCM2 phosphorylation (MCM2 pS53) versus DNA content in untreated cells and cells treated with 5 h auxin, 3 h CDC7 inhibitor (CDC7i), or 5 h auxin plus 3 h CDC7i. Pseudocolor plots as in (a). (l) Quantification of (k). Bars represent mean ± SD; statistical comparisons by two-way ANOVA with Tukey’s multiple comparisons test. (m) EdU incorporation versus DNA content under untreated, 5 h auxin, 3 h CDC7i, and 5 h auxin plus 3 h CDC7i conditions. Pseudocolor plots as in (a). (n) Quantification of (m), analyzed as in (g).

We next determined whether RIF1 loss stimulates replication by increasing origin initiation or by accelerating fork progression. To distinguish between these possibilities, we performed nanopore replication analysis after sequential EdU and BrdU labeling in the presence of auxin, 1NM-PP1, or both (Fig. 6H). RIF1 depletion increased fork progression while reducing origin firing (Fig. 6I, J). CDK1 inhibition alone reduced fork progression, whereas combined auxin and 1NM-PP1 treatment increased fork progression to levels comparable to RIF1 depletion alone (Fig. 6I). Initiation rates were not significantly altered by CDK1 inhibition (Fig. 6J). These results indicate that RIF1 loss bypasses CDK1 inhibition primarily at the level of fork progression, placing RIF1 downstream of CDK1 in this pathway.

RIF1 recruits PP1 phosphatase to reverse DDK-mediated MCM phosphorylation (65-67). Therefore, the canonical explanation for elevated DNA synthesis after RIF1 depletion is that more DDK-phosphorylated MCMs become available for origin firing. If this were the primary mechanism, then DDK inhibition should both reduce MCM2 phosphorylation and block the DNA synthesis increase caused by RIF1 loss. To test this, we measured phospho-MCM2 (Ser53) and EdU incorporation after RIF1 depletion, XL-413-mediated DDK inhibition, or both (Fig. 6K–N).

MCM2 phosphorylation changed upon DDK inhibition or RIF1 depletion as the canonical model predicts: XL-413 reduced phospho-MCM2, RIF1 depletion increased it, and XL-413 fully prevented that increase (Fig. 6K, L). DNA synthesis, however, did not follow the same pattern. With DDK inhibited, RIF1 depletion still restored EdU incorporation nearly to control levels (Fig. 6M, N), even though XL-413 had eliminated the RIF1-depletion-induced rise in MCM phosphorylation. Increased DDK-dependent MCM phosphorylation is therefore not required for the elevated DNA synthesis caused by RIF1 loss. Together with the nanopore data showing reduced initiation frequency after RIF1 depletion, these results indicate that RIF1 suppresses DNA synthesis primarily by restraining fork progression in these cells rather than by limiting origin activation. These findings place RIF1 downstream of CDK1 as a restraint on fork progression and indicate that CDK1-mediated RIF1 inhibition is the mechanism by which cells sustain DNA synthesis when origin firing is reduced.

## DISCUSSION

MTBP is a metazoan replication-initiation factor required for helicase activation and origin firing (17). Consistent with this established role, acute MTBP depletion strongly suppressed DNA synthesis and reduced CDC45 chromatin loading in HCT116 cells. Unexpectedly, however, DNA synthesis partially recovered over time despite persistent MTBP depletion. This recovery occurred without restoration of CDC45 loading or normal origin firing. Instead, our data support a model in which cells partially preserve replication output through CDK1-dependent increases in fork progression.

Both MTBP depletion and ATR inhibition increase fork progression through CDK1, though ATR inhibition produces this effect only when MTBP is depleted. Although CDK2 is generally regarded as the dominant S-phase CDK, prior studies have shown substantial functional overlap between CDK1 and CDK2, particularly under stress or checkpoint-perturbed conditions (37-39, 44, 45, 68-71). Under normal conditions, ATR inhibition activates CDK2 as well as CDK1 (44, 45, 56, 57).

Without MTBP knockdown ATR stimulated replication through a mechanism not dependent on CDK2. We propose that the resulting increase in CDK2 activity promotes origin firing and dominates the replication response, masking the CDK1-dependent contribution to fork rate. When MTBP is depleted, CDK2 cannot drive origin firing because MTBP is required for that step. This genetic separation isolates the CDK1/fork-rate arm of the ATR response. ATR inhibition in MTBP-depleted cells increased DNA synthesis without restoring CDC45 loading, and this effect required CDK1. Together, these results reveal an origin-firing-independent function of the ATR-CDK1 axis: ATR normally restrains fork speed during S phase by suppressing CDK1, and this restraint is relieved by both MTBP depletion and direct ATR inhibition.

Bulk DNA synthesis reflects the combined contributions of initiation frequency and fork progression, and prior studies have established that these two processes are inversely regulated (48, 59). Consistent with this relationship, nanopore replication analysis demonstrated reduced initiation together with increased fork progression during the adaptive state. Our findings suggest, however, that this shift is not merely a passive consequence of reduced origin usage. The genetic interactions among CDK1, ATR, and RIF1 indicate that fork progression is actively regulated through checkpoint-associated signaling. Earlier work showed that CDK1 activated during S phase stimulates replication by phosphorylating RIF1 (44, 45, 72), suggesting that CDK1-dependent inhibition of RIF1 accounts for the increased fork progression in MTBP-depleted cells.

Consistent with this model, RIF1 depletion phenocopied the increase in fork progression observed after MTBP depletion. RIF1 is best known as a PP1-targeting factor that opposes DDK-dependent MCM phosphorylation and suppresses origin activation (63, 65-67, 73-77). As expected, RIF1 depletion increased MCM phosphorylation and EdU incorporation in our system. However, nanopore replication analysis revealed that RIF1 depletion reduced initiation frequency while increasing fork progression. Thus, in this context, increased MCM phosphorylation was not sufficient to increase global origin firing, and faster forks rather than increased initiation best explained the elevated DNA synthesis after RIF1 loss. This finding is consistent with Yamazaki et al., who reported that RIF1 knockdown in HeLa cells increased replication track lengths together with inter-origin distance, indicating increased fork progression and reduced origin firing (64).

Similarly, work in Drosophila demonstrated that Rif1 can directly restrain fork progression during developmental gene amplification (78). Together, these observations indicate that RIF1 regulates fork progression in addition to its established role in origin firing.

The adaptive response partially restored replication output but was weakest in late S phase, where the effects of MTBP depletion were minimal compared with early and mid S phase. Rather than indicating a ceiling on fork-based compensation, this spatial pattern suggests that CDK1-driven fork acceleration is already at or near capacity in late S phase under basal conditions, leaving little room for additional increase. If so, the CDK1-RIF1 axis described here may reflect normal late-S-phase biology rather than a purely compensatory response to replication stress. As S phase advances, declining origin firing and attenuating ATR-mediated checkpoint signaling would permit rising CDK1 activity to accelerate forks (30, 33), progressively shifting cells toward a fork-progression-dominated mode of replication. This CDK1-dependent shift may represent a form of replication plasticity in which the balance between initiation frequency and fork rate adjusts dynamically over the course of S phase. Such a shift may be particularly important in late-replicating heterochromatic domains, where origin density may be low and completion of replication would depend in part on extended fork progression (79, 80). Whether this CDK1-dependent acceleration is adaptive, helping cells complete replication as origin firing declines, or whether sustained acceleration incurs replication stress cost (59, 81), remains to be determined.

## MATERIALS AND METHODS

### Cell culture and cell lines

HCT116-derived cell lines were cultured under the same conditions as described previously (53). Cells were maintained at 37°C in a humidified incubator with 5% CO_2_ and were routinely monitored for mycoplasma contamination. The MTBP degradation line used in this study was derived from the previously described HCT116 MTBP-mClover-miniAID line (53). To enable auxin-inducible degradation, this line was transduced with a retrovirus expressing Oryza sativa TIR1, generating HCT116 MTBP-mClover-miniAID; retroviral OsTIR1 cells, referred to here as MTBP-AID cells.

The HCT116 RIF1-AID2 cell line was obtained from Masato Kanemaki and was previously described in Klein et al. (62). This line is referred to here as RIF1-AID2 cells. CDK1 analog-sensitive derivatives of the MTBP-AID and RIF1-AID2 cell lines were generated using the one-shot strategy described previously (33). Briefly, cells were cotransfected with pCMV(CAT)T7-SB100, CDK1as_T2A_Zeo, and pX330_human CDK1, selected with zeocin, and screened for sensitivity to 1NM-PP1. A CDC45-mClover/CDK1AS cell line was generated using the same strategy from the previously described HCT116 CDC45-mClover background (41).

### Auxin-inducible degradation and drug treatments

Auxin-inducible degradation was performed by treating cells expressing mClover-miniAID-tagged MTBP with 500 μM indole-3-acetic acid (IAA; auxin; Sigma, I5148-2G) or AID2-tagged RIF1 with 1 μM 5-Phenyl-1H-indole-3-acetic acid (5-Ph-IAA; GLPBIO GC61525). Loss of MTBP or RIF1 was confirmed by flow cytometric measurement of total and chromatin-bound anti-GFP signal.

Drug treatments were performed by adding inhibitors directly to the culture medium for the times indicated in the figure legends. CDK1AS cells were treated with 5 μM 1NM-PP1 (Calbiochem, 529581) to inhibit analog-sensitive CDK1. ATR kinase was inhibited with 5 μM AZD6738 (ATRi; APExBIO, B6007 / 501152496), prepared by diluting a 5 mM stock 1:1000. CDC7/DDK was inhibited with 10 μM XL-413 (DDKi; Selleck Chemicals, S7547).

### siRNA transfection

siRNA transfections were performed as described previously (53). Briefly, cells were transfected with siRNAs using Lipofectamine RNAiMAX in Opti-MEM reduced-serum medium (Life Technologies), and cells were analyzed after the knockdown intervals indicated in the figure legends. MTBP was depleted using siGENOME Human MTBP SMARTpool siRNA (M-013953-01-0010; Dharmacon). CCNB1 was depleted using a pool of three AccuTarget genome-wide predesigned siRNAs targeting human CCNB1 (Bioneer; 891-1, 891-2, and 891-3). Negative-control siRNA was the same as described previously (53). Knockdown efficiency was assessed by flow cytometry using antibodies against the corresponding target protein or epitope tag.

### EdU incorporation and flow cytometry analysis of DNA synthesis

DNA synthesis was measured by EdU incorporation as described previously (41). Briefly, cells were pulse-labeled with 20 μM EdU for 30 min before collection. EdU incorporation was detected using Click-it chemistry, and DNA content was measured by propidium iodide staining. Samples were analyzed on a CytoFLEX LX flow cytometer (Beckman Coulter), and data were analyzed with FlowJo v10.10. Cells were gated by DNA content to define G1, early S, mid S, late S, and G2/M populations. Median EdU signal was quantified within the indicated cell-cycle gates and normalized as described in the corresponding figure legends.

### Total, nuclear, and chromatin-bound protein flow cytometry

Total, nuclear, and chromatin-bound protein levels were measured by flow cytometry using approaches adapted from previously described methods (53). For total protein analysis, cells were fixed and stained without detergent pre-extraction. For nuclear CCNB1 analysis, cells were processed to generate crude nuclear fractions before fixation. Briefly, cell pellets were resuspended in Buffer A (10mM Hepes, 10mM KCl, 1.5mM MgCl2, 340mM Sucrose, 10% Glycerol and 0.1% TritonX-100). Samples were incubated on ice, centrifuged to remove the cytosolic fraction, and the remaining nuclei were fixed with 4% paraformaldehyde in PBS. Nuclei were blocked in PBS containing 2.5% normal goat serum (NGS, Jackon Immunoresearch) and incubated with primary and secondary antibodies in PBS/NGS containing 0.1% NP-40. For chromatin-bound protein analysis, soluble proteins were first removed by CSK pre-extraction before fixation and antibody staining, leaving the detergent-resistant fraction. DNA content was measured by propidium iodide staining, and samples were analyzed on a CytoFLEX LX flow cytometer (Beckman Coulter). Data were analyzed with FlowJo v10.10.

mClover-tagged MTBP, RIF1, and CDC45 were detected using anti-GFP antibody (Rockland, 600-401-215S). CCNB1 was detected using mouse anti-Cyclin B antibody (BD Biosciences, 610219; lot 9100943). Phospho-histone H3 Ser10 was detected using the same antibody described previously (53). Phospho-MCM2 Ser53 was detected using anti-phospho-MCM2 Ser53 antibody (Bethyl/Fortis, A300-756A; lot 4). Alexa Fluor 488- or Alexa Fluor 647-conjugated secondary antibodies were obtained from Life Technologies.

Cells or nuclei were gated by DNA content to define G1, early S, mid S, late S, and G2/M populations. Gates were defined using the untreated or control sample and applied uniformly across matched treatment conditions. Background signal was determined using untagged cells or no-primary-antibody controls, as appropriate, and median fluorescence intensity was quantified within the indicated cell-cycle gates.

### Nanopore sequencing and replication-track analysis

Replication-track analysis by nanopore sequencing was performed as described previously (41). Briefly, cells were sequentially pulse-labeled with 50 μM EdU for 10 min followed by 50 μM BrdU for 10 min, then chased with thymidine for 40 min before collection. High-molecular-weight genomic DNA was isolated and processed for Oxford Nanopore sequencing using the same library-preparation and sequencing workflow described previously (41). Reads were base-called, aligned to the human hg38 reference genome, and analyzed with DNAscent and forkSense using the same parameters as described previously (41). Replication tracks were quantified from EdU- and BrdU-labeled reads. Fork progression was analyzed as total replication-track length. Origin usage was quantified as the percentage of initiation events relative to total forks. Treatment timing and normalization are described in the corresponding figure legends.

### Statistical analysis

Statistical analyses were performed as described in the corresponding figure legends. Simple linear regression was used where indicated. For comparisons across multiple conditions and cell-cycle fractions, two-way ANOVA followed by Tukey’s multiple-comparisons test was used unless otherwise specified. Data are presented as mean ± SD or mean ± SEM, as indicated in each figure legend. Replicate numbers are reported in the figure legends.

## Data availability

Flow cytometry data, FlowJo workspaces, nanopore sequencing data, and processed replication-track analysis files will be made available through public repositories upon publication.

## ACKNOWLEDGMENTS

We thank Masato Kanemaki and David Gilbert for the HCT116 RIF1-AID2 cell line. We thank the Bioinformatics and Pathways Core, supported by COBRE grant 5P30GM149376-02, for assistance with data analysis. Flow cytometry was performed at the OMRF Flow Cytometry Core, imaging was performed at the OMRF Imaging Core Facility, and nanopore sequencing was performed at the OMRF Clinical Genomics Center. Data processing and analysis were supported by the OMRF Center for Biomedical Data Sciences. The following plasmids were obtained from Addgene: pBabe Puro osTIR1-9Myc (#80074; Andrew Holland), pX330_human CDK1 (#118597; William Earnshaw), and CDK1as_T2A_Zeo (#118596; William Earnshaw). Funding provided by National Institutes of Health [R01GM157525] and The Presbyterian Health Foundation. MSH received support from the John and Mildred Carson PhD Scholarship Fund awarded for the OMRF Pre-doctoral Scholarship.

